# Rhamnolipid production from waste cooking oil using newly isolated halotolerant *Pseudomonas aeruginosa* M4

**DOI:** 10.1101/2020.03.09.983478

**Authors:** Juan Shi, Yichao Chen, Xiaofeng Liu, Yi Ran, Dong Li

## Abstract

This study isolated a novel halotolerant *Pseudomonas aeruginosa* M4, that was able to degrade oil and produce rhamnolipids. Various carbon sources, nitrogen sources, inoculum ratio, pH, and temperature were tested to optimize the oil degradation conditions. The highest oil degradation rate of 85.20 % and lipase activity of 23.86 U/mL were obtained under the optimal conditions (5% inoculum at 35 °C and pH 8). The components of degradation products at different times were analyzed to explore the mechanism of oil degradation by GC-MS. Short chain fatty acid of acetic and n-butyric acids were the primary degradation intermediates. *P. aeruginosa* M4 had good salt tolerance up to 70 g/L. The maximum rhamnolipid concentration of 1119.87 mg/L was produced when *P. aeruginosa* M4 used waste cooking oil as the sole carbon source. Rhamnose precursors were synthesized from glycerol, a hydrolysis product of waste cooking oil. R-3-hydroxyalkanoate precursors were synthesized de novo using acetyl-CoA produced from β-oxidation of fatty acids. The findings show that *P. aeruginosa* M4 is a valuable biosurfactant producer in the treatment of waste cooking oil.

**Key Points:** *P. aeruginosa* isolation, oil degradation mechanism, rhamnolipid production from WCO

## Introduction

Food waste (FW) grows steadily every year. Untreated FW causes serious environmental problems, including soil pollution, odor production, and pest attraction. Therefore, FW management is an urgent need worldwide, with a growing emphasis on resource recovery and environmental protection promoting the rapid adoption of novel technologies in this field (Zhang et al. 2010; HKEPD 2015). The composition of FW is highly heterogeneous, but mostly includes (w/w) 40-60% starch, 5-10% protein, 0.84-1.38% salt, and 10-40% fatty or oily contents (Pleissner et al. 2013; Demichelis et al. 2017). In general, fatty or oily waste products, specifically waste cooking oil (WCO), can be separated and collected from the FW (Patil et al. 2012).

FW management currently includes landfilling, incineration, and biological treatments like composting, anaerobic digestion, or transformation to animal feed (Wang et al. 2013; Kiran et al. 2015; Chan et al. 2016; Pleissner et al. 2017). Landfilling and incineration are rarely used because of the high water content of FW, and when FW is used as animal feed, its ingredients are transferred into the food chain, affecting the entire ecosystem and ultimately influencing human health (Chen et al. 2009; Lan et al. 2015). Biological treatment of FW is challenging because of the large amounts of oil in it. In many cases, an oil layer will form on the surface of the FW preventing the diffusion of oxygen, which reduces composting efficiencies. Oil also poses a barrier to anaerobic digestion because of its inhibitory effects on anaerobic microorganisms and high rates of digester foaming (Awasthi et al. 2018; Cammarota and Freire 2006).

One of the most economic and environmentally friendly approaches to dealing with FW was to use the WCO to produce biosurfactants (Sharma et al. 2019; Maddikeri et al. 2015; Thavasi et al. 2007). Biosurfactants can emulsify hydrophobic hydrocarbons, enhancing their solubility in water and reducing surface tension. In addition, biosurfactants have much lower toxicity and higher biodegradability, structural diversity, and surface activity than chemical surfactants (Dobler et al. 2016). However, few studies have focused on biosurfactant production using WCO as the sole carbon source, probably because of differences in the quality of WCO from different sources (Pansiripat et al. 2010). To date, most reports on biosurfactant production from WCO have been less than remarkable with maximum oil degradation achieved almost a week later (Gudina et al. 2013; Rabiei et al. 2013).

The purpose of this study was to isolate a highly efficient oil degrading strain that can produce biosurfactants. Soybean oil was used as the limiting carbon source to evaluate the oil degrading ability of strains. The conditions for oil degradation (carbon source, nitrogen source, inoculum, temperature, and initial pH) were optimized. GC-MS was used to analyze the degradation products at various time points to explore the mechanism of oil degradation. The ability of the strain to produce biosurfactants and resist salt stress was also investigated using WCO as the sole carbon source.

## Materials and methods

### Samples and culture medium

Soil samples were collected from an FW treatment plant and used to isolate target microorganisms. Nutrient broth (NB) (seed medium) contained beef extract 5.0 g/L, tryptone 10.0 g/L, and NaCl 5.0 g/L. For the preparation of nutrient agar plates and slants, 15 g/L agar (strength 1300) was also added to the NB. Mineral salt medium (MSM) contained (NH_4_)_2_SO_4_ 5.0 g/L, K_2_HPO_4_ 2.0 g/L, KH_2_PO_4_ 2.0 g/L, MgSO_4_ 0.5 g/L, NaCl 2.0 g/L, 2 mL trace element solution (ZnSO_4_·7H_2_O 0.66 g, MnCl_2_·4H_2_O 0.44 g, H_3_BO_4_ 0.02 g, CoCl_2_·6H_2_O 0.40 g, CuCl_2_·2H_2_O 0.01 g, NiCl_2_·6H_2_O 0.61 g, NaMoO_4_·H_2_O 0.73 g, EDTA 0.01 g, and 1 L deionized water). Oil degradation medium was prepared by adding soybean oil to 1 L MSM. Rhamnolipid production broth comprised WCO and 1 L MSM. The pH of all the media was adjusted to 7 using 1 N NaOH and 1 N HCl and autoclaved at 121 °C for 20 min before use.

The fatty acids in soybean oil used in this study primarily contains 1984 mg/L and 8848 mg/L of palmitic and linoleic acids, respectively.

### Isolation and identification of oil degrading strains

The isolation and identification protocols were modified from those described by Zhang et al (2012). Briefly, about 10 g of each soil sample was added to a flask containing sterile glass beads and sterile water. Following mixing, 10 mL of the supernatant was inoculated into enriched culture medium and incubated at 35 °C and 180 rpm until the oil was degraded. Thereafter, the oil concentration in the medium was gradually increased (5, 10, 15, 20 and 25 g/L) every 6 days. One milliliter of the last acclimated bacterial solution was diluted in a gradient, and the 10^-5^, 10^-6^, and 10^-7^ bacterial solutions were spread on lipase screening plates and inverted in a biochemical incubator for 1-3 days. Large and bright red colonies were then picked and inoculated in seed medium. The seed broth was then inoculated at 5% (v/v) into re-screening medium with soybean oil as the only carbon source and cultured for 3 days. The residual oil, lipase activity, and cell density of the fermentation broth were measured to evaluate the oil degrading ability of the isolated strains. The strain with the highest soybean oil degradation ability was then selected and incubated on agar slant medium, cultured at 35 °C for 24 h, and stored at 4 °C for the next set of evaluations. This strain was then identified using phylogenetic analysis, physiological and biochemical identification, and 16S rDNA sequencing. Morphological characteristics were evaluated using a scanning electron microscope. 16S rDNA sequence analysis was performed using universal primers 27F (seq) and 1492R (seq). These sequences were then compared with known 16S rDNA sequences using BLAST, and phylogenetic analysis was constructed using the Neighbor-Joining method in MEGA version 6.0 software (Dastager et al. 2009).

### Optimization of oil degradation conditions

We used a systematic approach to evaluate all the nutrient and physical parameters in the oil degradation assays (Kantak et al. 2011). All selected strain inoculums were prepared from NB medium after incubation at 35 °C for 20 h. The optimization experiments were all conducted in 150 mL Erlenmeyer flasks with 30 mL culture medium at 35 °C and 180 rpm. In this analysis, we evaluated two nutrient and three physical parameters to create the optimal culture conditions for oil degradation. These parameters included carbon source (soybean oil, glucose, fructose, malt extract powder, sucrose, and soluble starch), nitrogen source ((NH_4_)_2_SO_4_, NaNO_3_, urea, tryptone, and yeast extract), inoculum ratio (2–10%, v/v), temperature (25–60 °C), and pH (4-10). After 3 days of fermentation, the cell density in the fermentation broths was measured, and the lipase activity of the fermentation supernatant was determined. The fermentation supernatant was obtained by centrifuging the fermentation broth at 8000 rpm for 10 min to remove bacterial cells and residual oil.

The oil degradation characteristics of the selected stain was evaluated after 5 days of fermentation carried out at the optimized conditions. Both cell growth and pH of the fermentation broth were observed during this time frame. Cell growth was monitored by optical density at 600 nm, and pH was recorded using a pH meter at 24 h intervals. The oil degradation rate and lipase activity of the fermentation broth were also measured every 24 h. Then, cell density, pH value, oil degradation rate, and lipase activity were plotted against time.

### Kinetics of rhamnolipid production

To study the kinetics of rhamnolipid production, the selected strain was inoculated into the fermentation medium and cultivated at 35 °C and 180 rpm. WCO was the only carbon and energy source in this fermentation medium, and its concentration was 5 g/L, 10 g/L, and 25 g/L. Samples were harvested every 24 h until reaching 132 h and analyzed for cell density, lipase activity, rhamnolipid concentration, and pH.

### Salt tolerance assessment

Seed solution with the selected strain was inoculated into the medium with a salt concentration of 0, 2, 5, 10, 15, 20, 25, 30, 50, 70, 100, and 150 g/L. In this experiment, uninoculated culture medium was used as a control, and the OD_600_ value of the culture medium was measured over time. The range of salt concentrations tolerated by the experimental strain was determined by the change in OD_600_ value.

## Analytical methods

### Measurement of cell density

Optical density was used to approximate cell density. A small amount of the fermentation broth was taken and divided into two groups, one which was centrifuged at 8000 rpm for 10 min, and the supernatant collected and used as the control, and the other was measured as the sample. The sample and the supernatant were diluted with sterile water to facilitate OD reading, and their optical density at 600 nm (OD_600_) was determined by a spectrophotometer. The cell density was obtained by subtracting the optical density of the supernatant from the optical density of the sample.

### Determination of lipase activity

Lipase activity was determined spectrophotometrically by measuring incremental changes in the absorption at 405 nm promoted by the hydrolysis of p-NPC (Reddy et al. 2016). The reaction mixture was composed of 0.7 mL of 0.05 M phosphate buffer (pH 8.0), 0.1 mL of p-NPC substrate solution, and 0.2 mL of diluted enzyme sample. It was incubated at 35 °C for 5 min, and then 1 mL of ethanol was added to terminate the reaction. The absorbance of resulting yellow product was measured at 405 nm. The substrate solution containing 32 μL p-NPC and 10 mL 2-propanol was prepared fresh for each round of analysis. One international unit of lipase activity was defined as the amount of enzyme needed to catalyze the release of 1 μmol of p-NP per minute from p-NPC under these conditions (Pacheco et al. 2012).

### Determination of oil degradation rate

Residual cooking oil content in the fermentation broth after incubation was quantified by infrared spectrophotometry. In accordance with the national standard for the determination of animal and vegetable oils, the remaining oil in the fermentation medium was extracted using an equal volume of tetrachloromethane. After mixing, the solution formed a stable upper aqueous phase and lower organic phase, and the lower organic phase was collected and diluted with tetrachloromethane. Then, 10 mL of this was taken and analyzed by an infrared spectrometer.

The oil content extracted from the control group was used as the initial oil concentration. The oil degradation rate was calculated by subtracting the oil content extracted from the experimental group from the initial oil content. This calculation can be represented by the following formula (1):

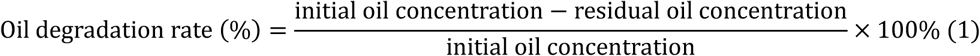

### Analysis of fatty acids produced during oil degradation

The supernatants from the fermentation broths were collected and freeze-dried at various time points (0, 12, 24, 36, 48, 72, and 96 h). These samples were sent to the Shanghai Minxin Biotechnology Co., Ltd. for fatty acid composition and content evaluation. The pretreatment for the determination of short-chain fatty acid content included the following steps: A sample of 300 μL was placed in a 1.5 mL centrifuge tube, then 115 μL sulfuric acid (50%) and 450 μL ether (20 μg/mL) containing an internal standard were added. This solution was mixed by vortex for 2 min and then placed into a centrifuge at 4 °C and run at 12,000 rpm for 20 min. Tubes were then placed at 4 °C for 30 min, and the supernatant was draw and used for GC-MS analysis (Radzuan et al. 2017). The pretreatment method for the determination of medium-long-chain fatty acids included the following steps: 1 mL of sample was added to a 10 mL centrifuge tube, then 2 mL of 10% acetylchloromethanol solution and 1 mL n-hexane were added, and the mixture was reacted at 95 °C for 2 h with oscillation every 5 min. After removal, it was cooled to room temperature, and 6 mL of 6% potassium carbonate was added. After centrifugation at 4000 rpm for 10 min, the supernatant was placed in a new 1.5 mL centrifuge tube and nitrogen was blown. 100 μL of n-hexane and 5 μL of the internal standard were added prior to redissolution. After centrifugation at 3000 rpm for 5 min, 100 μL of the supernatant was taken for evaluation.

### Rhamnolipid determination

The culture medium was first centrifuged at 8000 rpm for 10 min to remove cells for the quantitative analysis of rhamnolipid content. Five milliliters of the supernatant were then extracted with 15 mL of ethyl acetate. The upper organic phase was collected and evaporated until dry, and the rhamnolipid residues were dissolved in 2 mL of water. Rhamnolipid concentration in the cultivation medium was measured using the anthrone-sulfuric acid colorimetric method at 625 nm (Zhu et al. 2012), and the rhamnose value was calculated according to the L-rhamnose (20–100 mg/L) standard curve. The concentration of rhamnolipids was determined by multiplying the rhamnose value by a coefficient of 3.4, which was derived from the correlation of pure L-rhamnose (Benincasa et al. 2002). The anthrone-sulfuric acid solution was prepared by dissolving 0.1 g anthrone in 50 mL concentrated sulfuric acid solution.

## Results

### Isolation and characterization of oil degradation strains

Twenty-two bright red colonies were observed on the lipase screening plates after incubation at 35 °C for 1-3 days. After purification, the colonies were inoculated into the re-screening medium with soybean oil as the sole carbon source and incubated at 35 °C for three days. Only five strains were shown to grow well in the re-screening medium (Table 1). Strain M4 was chosen for further analysis as it had the highest growth rate, lipase activity, and oil degradation rate.

**Table 1.** Oil degradation characteristics of the five screened strains.

| Strain number | M4 | M8 | M9 | M12 | M14 |
| --- | --- | --- | --- | --- | --- |
| Cell density (OD<sub>600</sub>) | 2.81±0.13 | 2.12±0.02 | 2.67±0.08 | 2.51±0.02 | 2.11±0.06 |
| Lipase activity (U/mL) | 7.03±0.02 | 1.56±0.02 | 1.41±0.04 | 0.65±0.01 | 1.06±0.01 |
| Oil degradation rate (%) | 93.49±0.12 | 58.95±0.89 | 52.61±0.76 | 43.22±0.10 | 37.15±0.34 |

Physiological identification showed that M4 could grow well in a temperature range between 25 and 40 °C, with an optimum temperature at 35 °C; pH range was determined to be between 5 and 9, and the optimum value was 6. The morphological characteristic of M4 indicated by scanning electron microscope was shown in Fig. 1a, where M4 is a rod-shaped bacillus with a length of about 2.44 μm and a width of about 0.78 μm. A phylogenetic tree was constructed based on the 16S rDNA sequence (1400 bp) analysis using the Neighbor-Joining method (Fig. 1b). This analysis suggests that M4 is a strain of *Pseudomonas aeruginosa*, and BLAST analysis showed a 99% sequence similarity between this strain and *P. aeruginosa* DSM 50071.

**Fig. 1.**
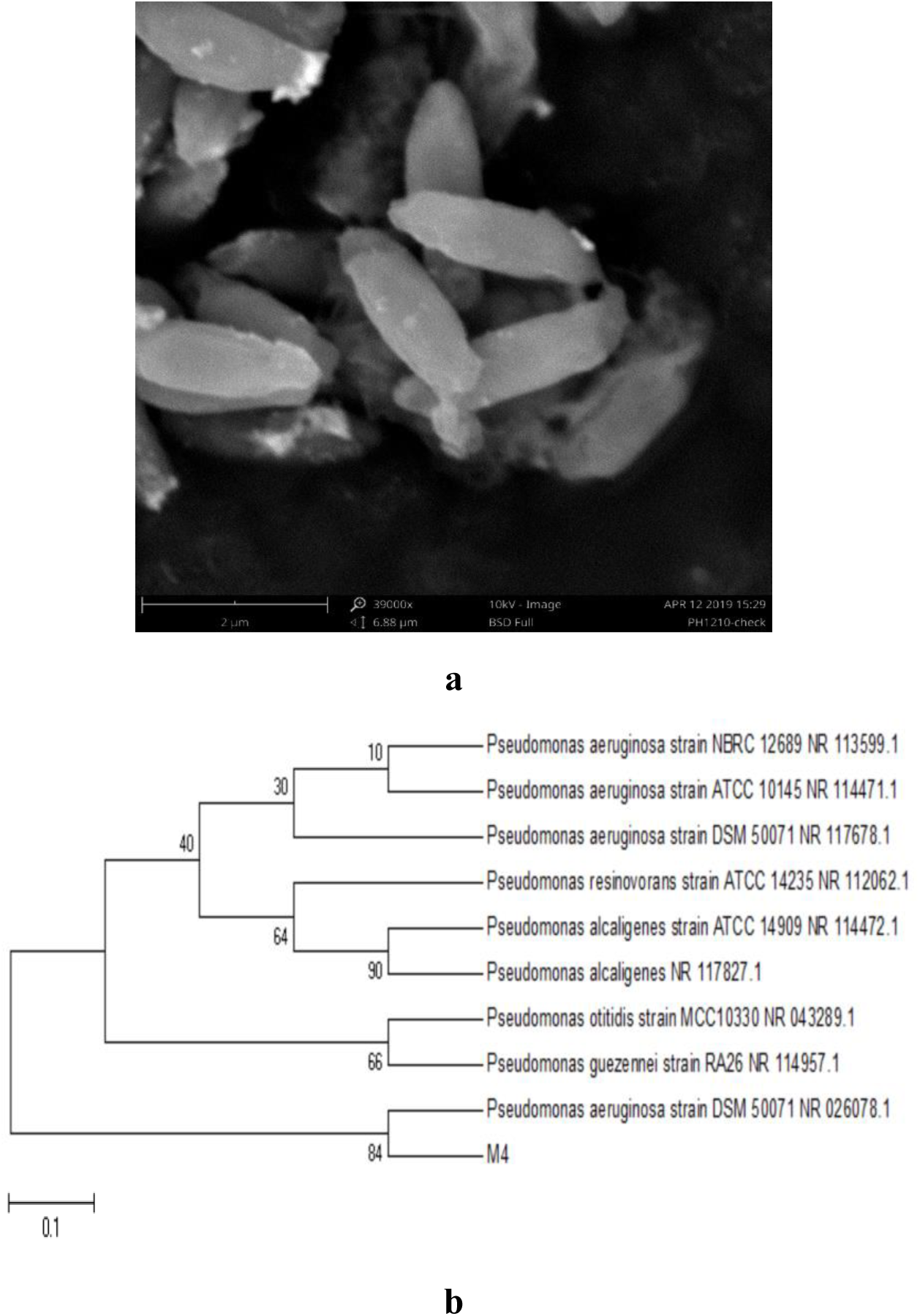
**a** Morphology of *P. aeruginosa* M4 shown by SEM. **b** Phylogenetic tree of *P. aeruginosa* M4 constructed by Neighbor-Joining algorithim.

### Optimization of oil degradation conditions for *P. aeruginosa* M4

#### Carbon and nitrogen sources

Different carbon sources can influence strain growth and lipase production. In this study, *P. aeruginosa* M4 was shown to have maximum lipase activity (11.20 U/mL) when soybean oil was available as a primary carbon source (Fig. 2a). Malt extract produced the highest biomass for *P. aeruginosa* M4, but despite this high biomass, lipase activity was relatively low, suggesting that in *P. aeruginosa* M4, lipase is inducible under specific growth conditions.

**Fig. 2.**
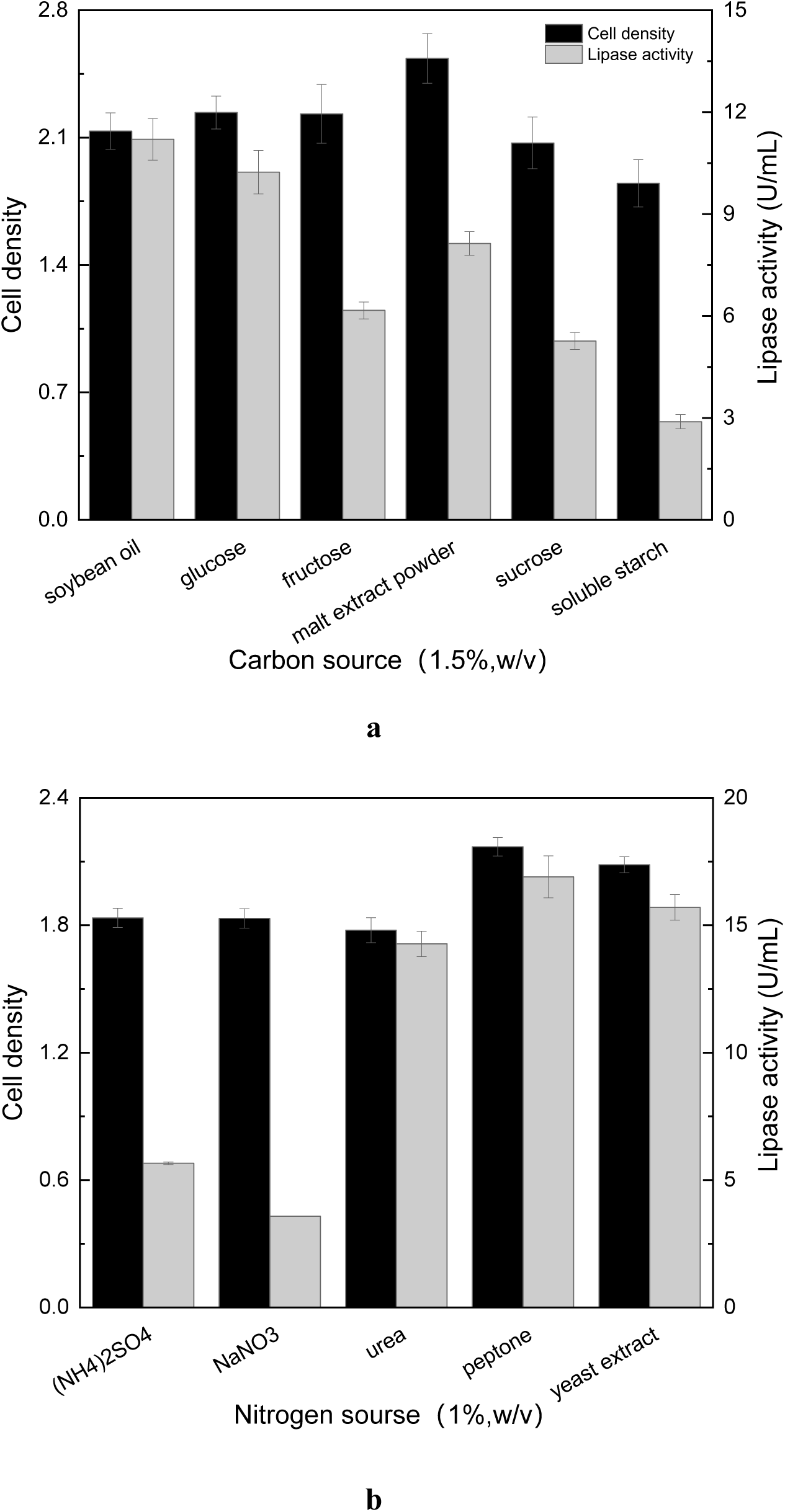
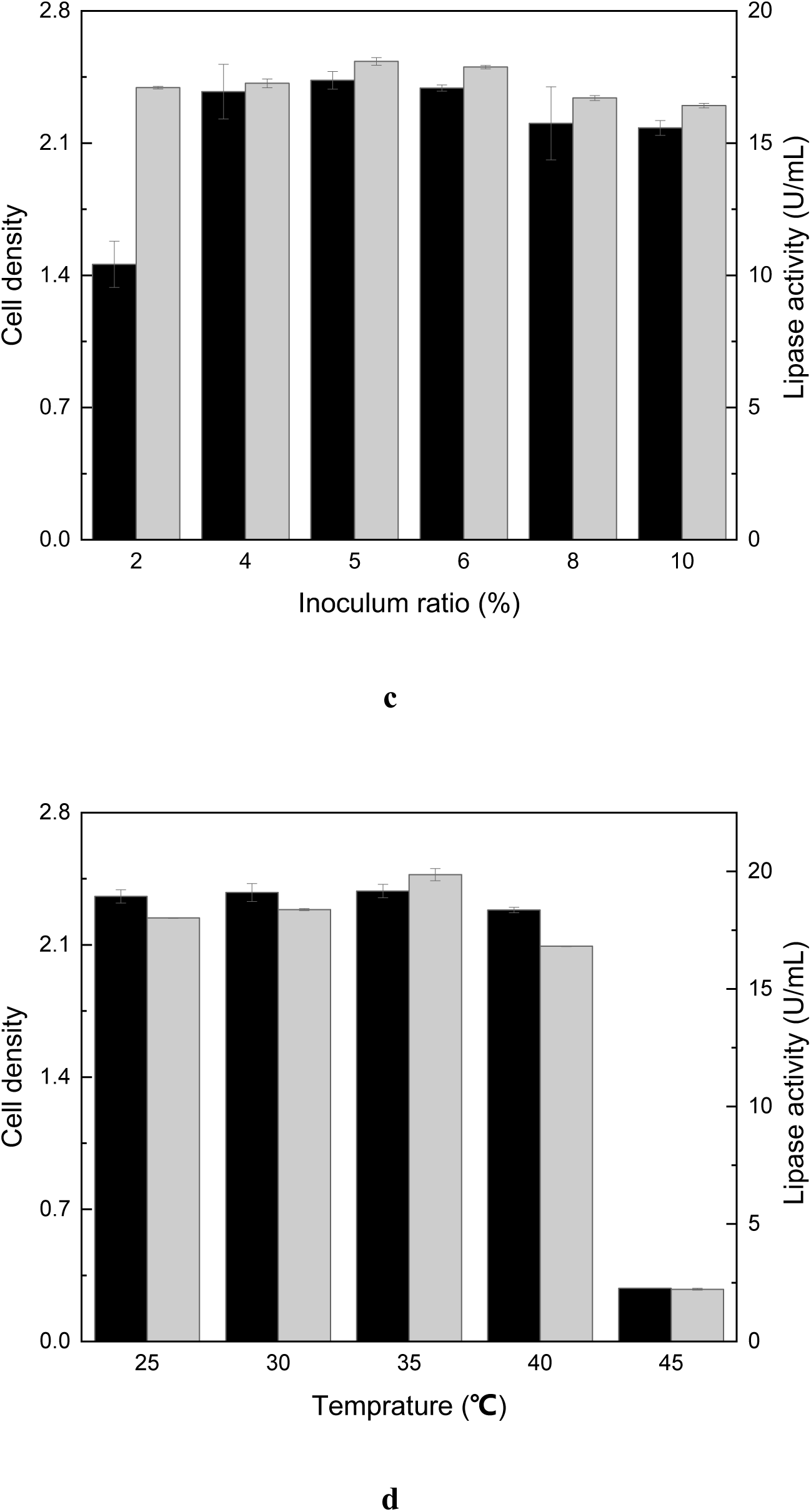
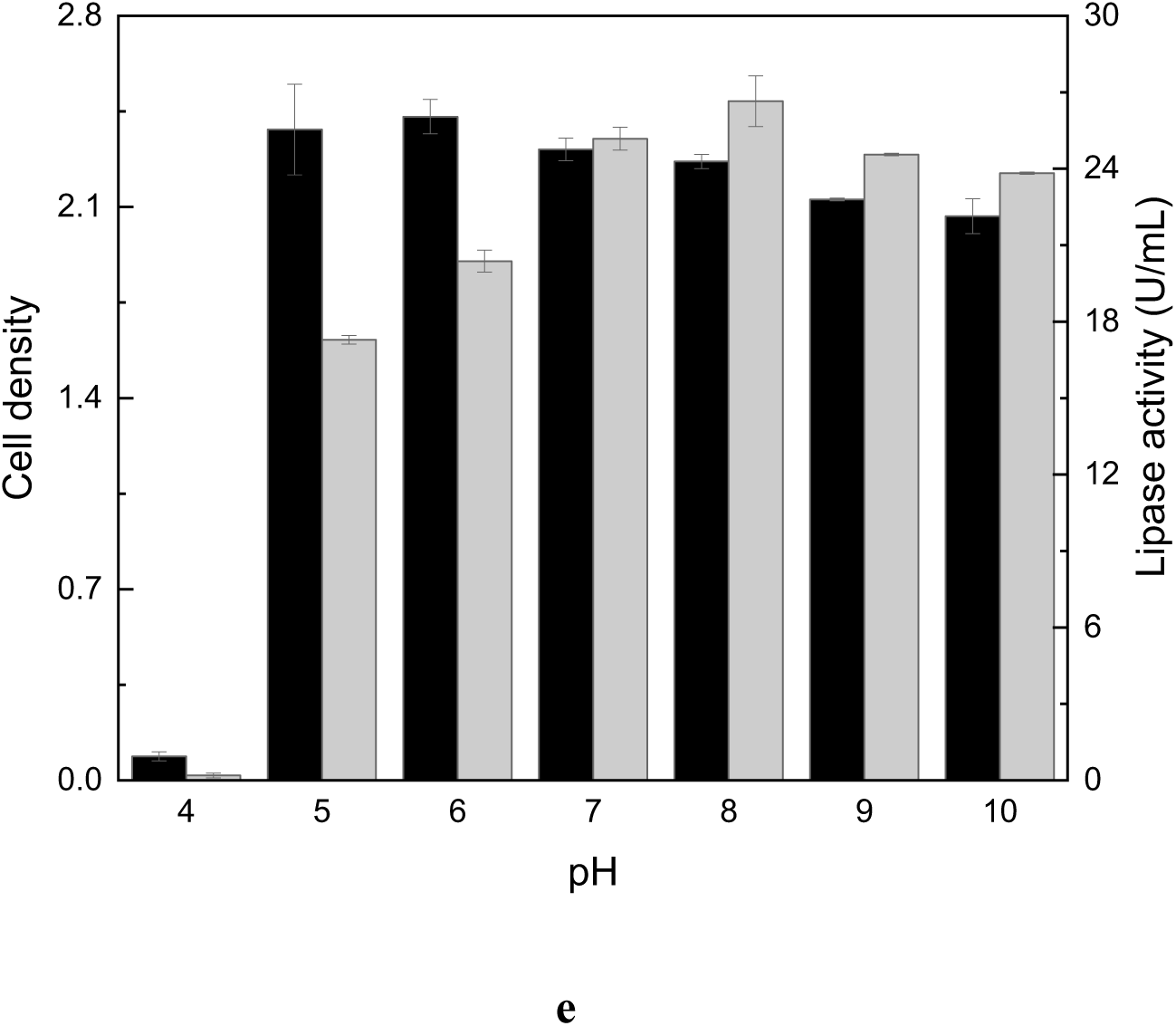
Change in cell density and lipase activity at various **a** carbon sources, **b** nitrogen sources, various **c** inoculum ratio, **d** temperature and **e** pH.

The various nitrogen sources tested in this study ((NH_4_)_2_SO_4_, NaNO_3_, urea, tryptone and yeast extract), the lipase activity and cell growth levels were highest (16.90 U/mL and 2.17, respectively) when tryptone was added to the fermentation medium (Fig. 2b). *Burkholderia spp.* displays a similar preference for tryptone when producing lipases (Gupta et al. 2007).

#### Inoculum ratio, temperature, and pH

The effect of the inoculum ratio on lipase activity was shown in Fig. 2c. The optimal inoculum value was shown to be 5% (v/v), as we observed significant growth inhibition when the inoculation ratio rose. Therefore, an inoculum ratio of 5% (v/v) was used in all the evaluations in following studies.

The lipase activity at different temperatures (25–45 °C) was determined. The highest lipase activity (19.87 U/mL) was obtained at 35 °C (Fig. 2d). But lipase activity steadily decreased as the temperature approached 45 °C.

The effect of pH was caused by protonation or deprotonation of participating molecules in one or more reaction steps (Taylor et al. 2002). Changes in the pH of the medium resulted in changes in lipase activity with the best activity (26.65 U/mL) observed at pH 8 and the worst activity at pH 4 (Fig. 2e). Previous studies have shown that *P. aeruginosa* and *Bacillus licheniformis* had the highest lipase activity at a pH between 7.0 and 10.0 (Mobarak et al. 2011; Sharma et al. 2011).

### Potential mechanism of oil degradation by *P. aeruginosa* M4

*P. aeruginosa* M4 was continuously cultured under optimum conditions in a fermentation medium supplemented with 20 g/L soybean oil as the sole carbon and energy source. The oil degradation rate, cell density, lipase activity, and pH of these fermentation broths are shown in Fig. 3. The oil degradation rate, cell density, and lipase activity increased with longer fermentation times. This was consistent with a previous report (Feng et al. 2017) of a similar trend. The decrease in pH over time may be related to the degradation of soybean oil and the production of fatty acids. During the 96 h fermentation, the oil degradation rate reached more than 85.20%, the cell density at OD_600_ was 2.32, and the lipase activity was 23.86 U/mL. The pH of the fermentation broth was lower than 5.3. With longer fermentation time, the accumulation of fatty acids and acidification of the fermentation broth negatively affected the growth and lipase activity of the strain, and the oil degradation rate was also reduced, suggesting that 96 h fermentation was the best time for oil degradation.

**Fig. 3.**
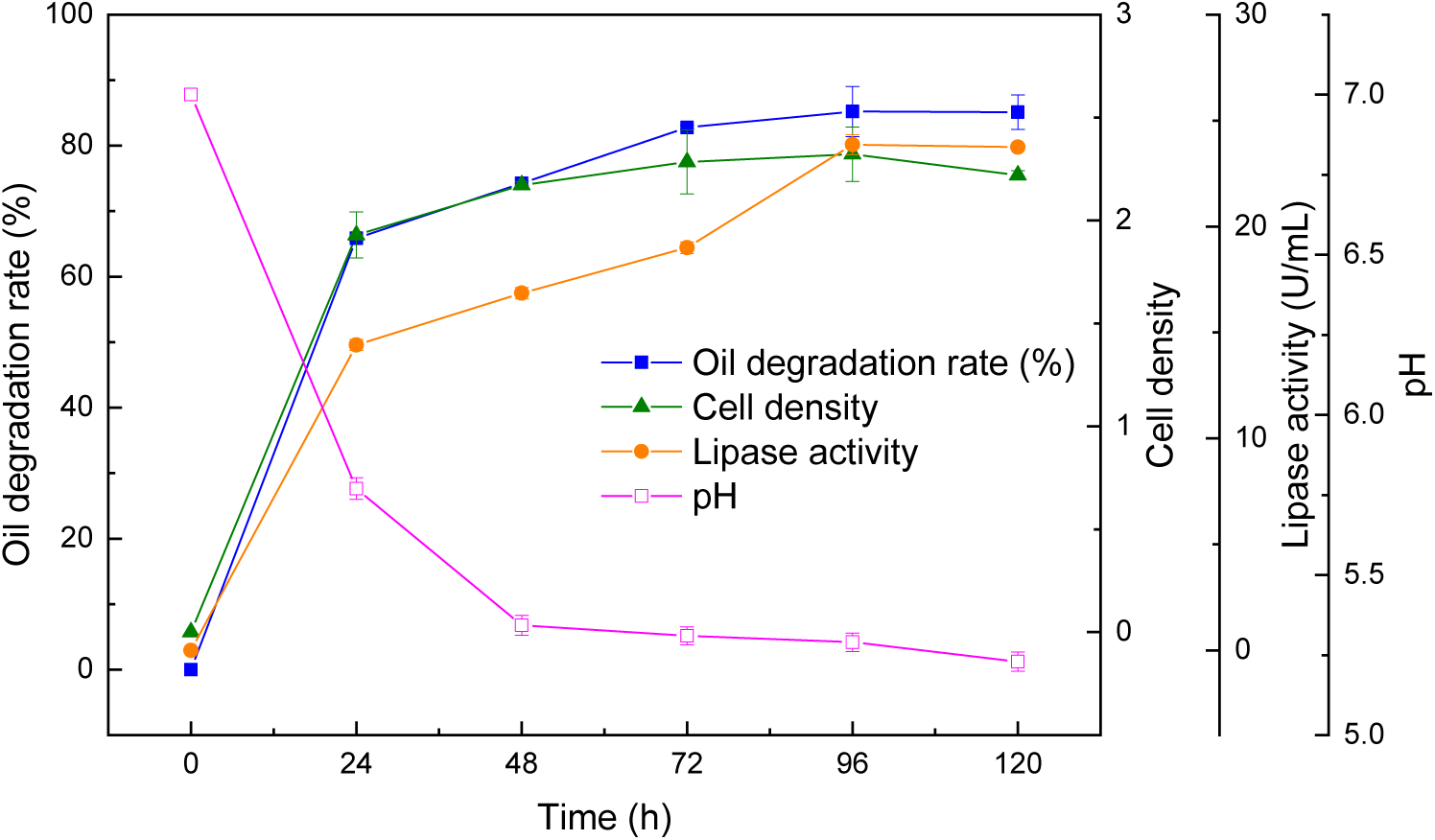
Oil degradation characteristics of *P. aeruginosa* M4 under optimal conditions.

Tables 2 and 3 show the medium-long-chain fatty acids and short-chain fatty acids identified in *P. aeruginosa* M4 degradation products at various time points. The main medium-long-chain fatty acids were C18 linoleic acid and C16 palmitic acid. Acetic and n-butyric acids were the main short-chain fatty acids. Vegetable oil contains saturated and unsaturated fats with a mixture of triglycerides, diglycerides, monoglycerides, and free fatty acids. In the process of metabolizing glycerides as carbon sources, they are first broken down by lipase into glycerol and fatty acids. Ramachandran et al. (2007) reported that most fatty acids, including palmitic and linoleic acids, found in vegetable oil are substrates for lipase hydrolysis. It is well known that the formation of acetic acid is through β-oxidation of long-chain-fatty acids. Butyric acid could be a degradation product of medium-long-chain fatty acids degradation or formed via the polymerization of these acetic acids. This pathway needs further analysis to understand its implications for fermentation.

**Table 2.** The content of medium-long chain fatty acids in the degradation products.

| **Fatty acid($\mu\text{g/mL}$)** | **12h** | **24h** | **36h** | **48h** | **72h** | **96h** |
| --- | --- | --- | --- | --- | --- | --- |
| **hexanoic acid(C6)** | 0.29 | 0.28 | 0.18 | 0.20 | 0.21 | 0.26 |
| **octanoic acid(C8)** | 0.17 | 0.69 | 3.15 | 13.71 | 7.75 | 6.96 |
| **decanoic acid(C10)** | 0.12 | 0.76 | 1.58 | 3.08 | 2.16 | 1.72 |
| **undecanoic acid(C11)** | 0.10 | 0.70 | 0.52 | 0.66 | 10.92 | 0.69 |
| **Lauric acid(C12)** | 0.51 | 2.67 | 3.64 | 9.64 | 8.36 | 14.91 |
| **tridecanoic acid(C13)** | 0.65 | 1.22 | 0.80 | 1.27 | 1.60 | 1.05 |
| **Myristic acid(C14)** | 2.90 | 4.21 | 4.49 | 6.96 | 9.75 | 7.32 |
| **pentadecanoic acid(C15)** | 3.90 | 1.37 | 1.68 | 2.91 | 2.94 | 3.02 |
| **Palmitic acid(C16)** | 36.96 | 109.85 | 143.71 | 336.23 | 600.28 | 508.61 |
| **heptadecanoic acid(C17)** | 16.69 | 25.69 | 7.60 | 12.44 | 17.63 | 12.43 |
| **linoleic acid(C18)** | 78.05 | 526.99 | 789.04 | 2118.99 | 5028.33 | 3637.10 |
| **arachidic acid(C20)** | 7.32 | 12.15 | 41.43 | 133.39 | 217.77 | 149.44 |
| **Henecanoic acid(C21)** | 16.10 | 1.66 | 2.33 | 5.47 | 6.10 | 5.73 |
| **behenic acid(C22)** | 43.77 | 18.55 | 39.44 | 74.93 | 102.49 | 92.55 |
| **tricosanoic acid(C23)** | 2.42 | 1.24 | 1.90 | 5.75 | 5.89 | 3.86 |
| **lignoceric acid(C24)** | 29.90 | 6.75 | 6.10 | 21.80 | 21.03 | 17.96 |
| **Total** | 239.87 | 714.77 | 1047.61 | 2747.43 | 6043.22 | 4463.61 |

**Table 3.**
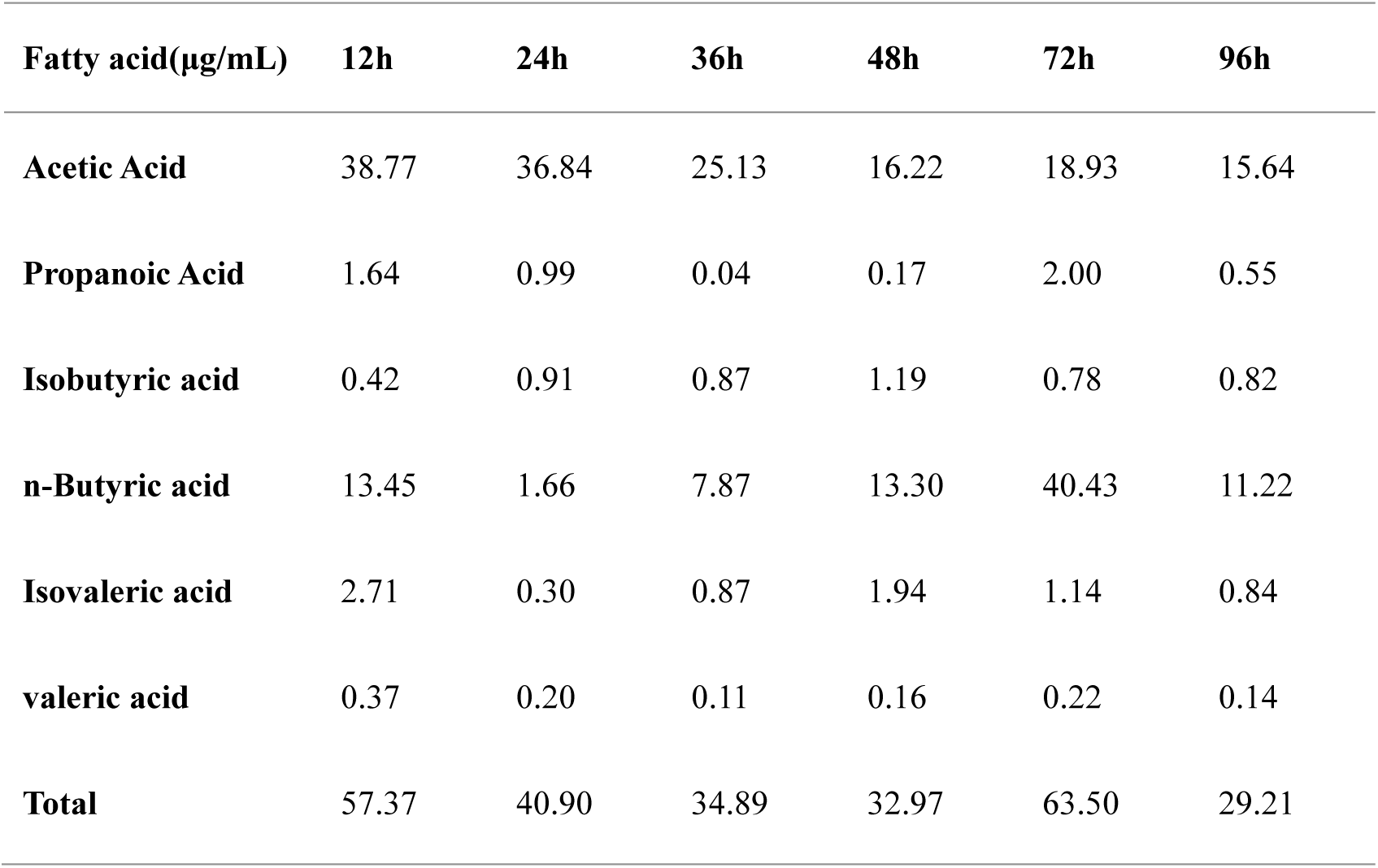
The contents of short chain fatty acids in the degradation products.

Acetic acid is used in cell biomass synthesis via the acetyl coenzyme A pathway, which is why we were able to observe very little accumulation of acetic acid during fermentation experiments. However, along with the hydrolyzation of the ester bonds by lipase, the total medium-long-chain fatty acid content increased and reached a maximum at 72 h. This shows that the hydrolysis rate is higher than the β oxidation rate in fermentations. From 72 h to 96 h, the concentrations of linoleic and palmitic acid decreased, which implies that the hydrolysis was completed at 72 h while the β oxidation continued up to the experimental end point.

As shown in Fig. 4, the clear medium turned turbid following fermentation, and the soybean oil on the surface of the medium had disappeared. This demonstrated that there was emulsification of the oil apart from degradation described above. It suggested there might be some biosurfactant produced as a byproduct in oil degradation process, which was confirmed in next part of the study.

**Fig. 4.**
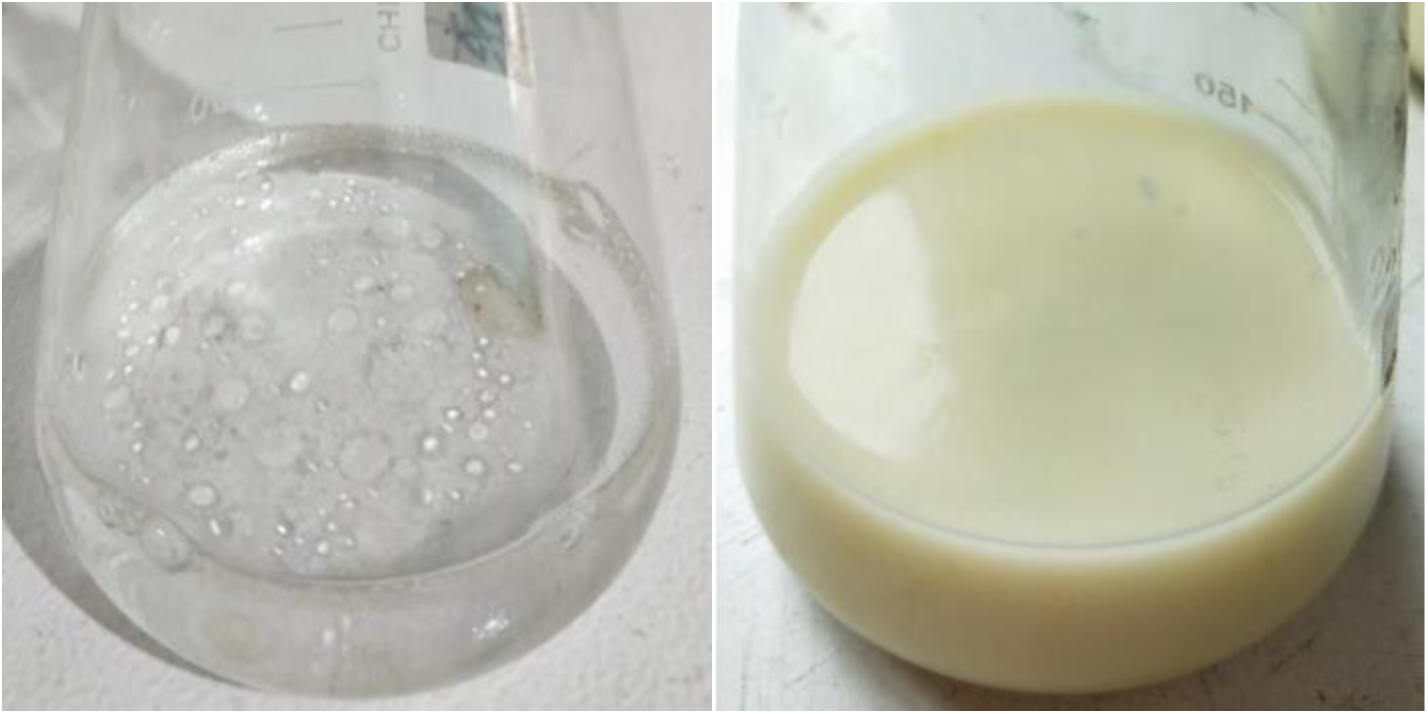
Photograph of degradation by *P. aeruginosa* M4 with soybean oil as the only carbon source.

### WCO degradation and rhamnolipid production by *P. aeruginosa* M4

The oil degradation and biosurfactant production were investigated using WCO as the only substrate (carbon and energy source). The fermentation broth was also shown to be emulsified, which suggests the presence and production of an emulsifying agent. Previous reports have suggested that other *P. aeruginosa* strains can produce biosurfactants belonging to the rhamnolipid family (Sharma et al. 2019). Thus, the biosurfactant analysis focused on this family of emulsifiers. The cell density, lipase activity, rhamnolipid concentration, and pH from the fermentations are shown in Fig. 5. The highest cell density of 2.61, the maximum lipase activity of 15.87U/mL, and the highest rhamnolipid concentration of 1119.87 mg/L were to be found.

**Fig. 5.**
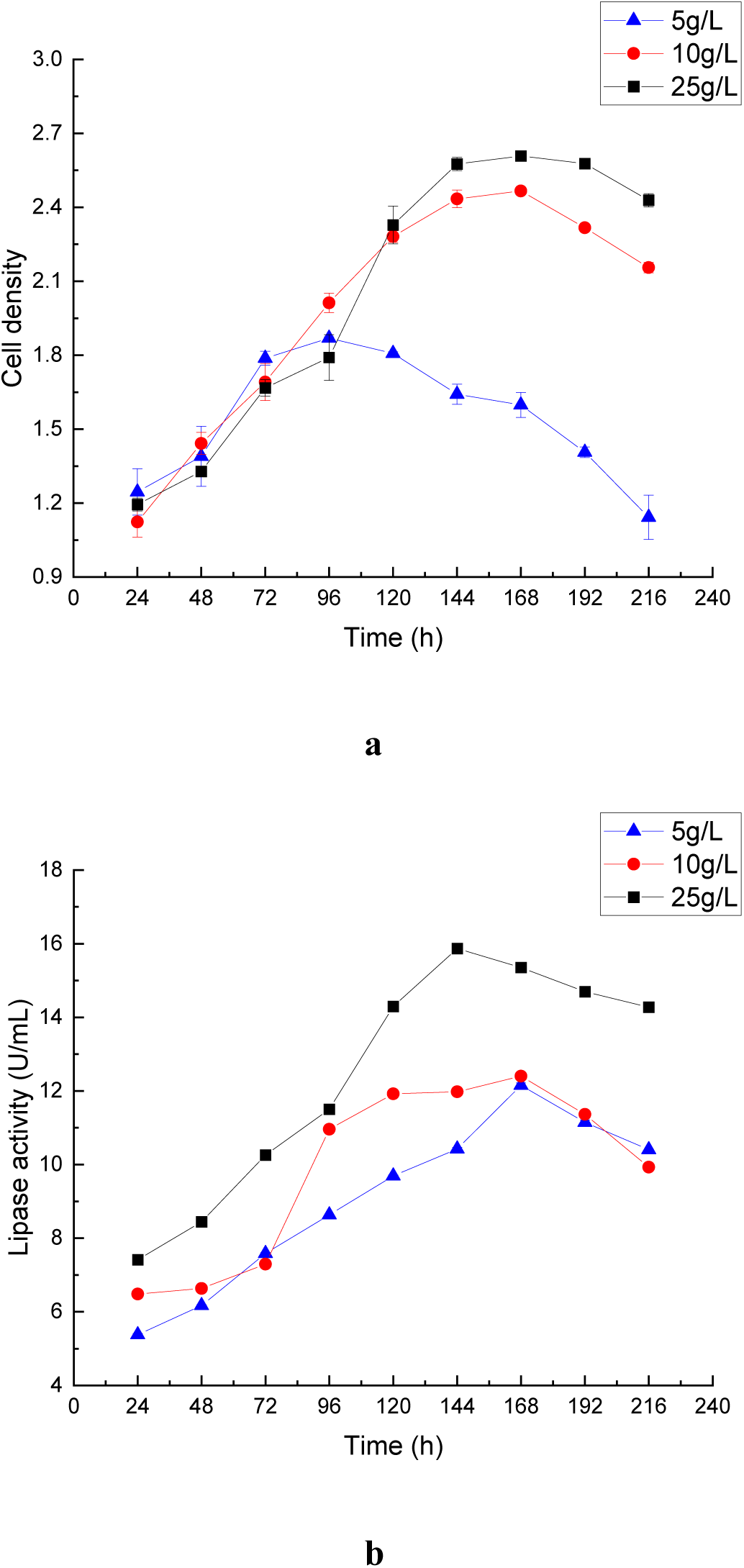
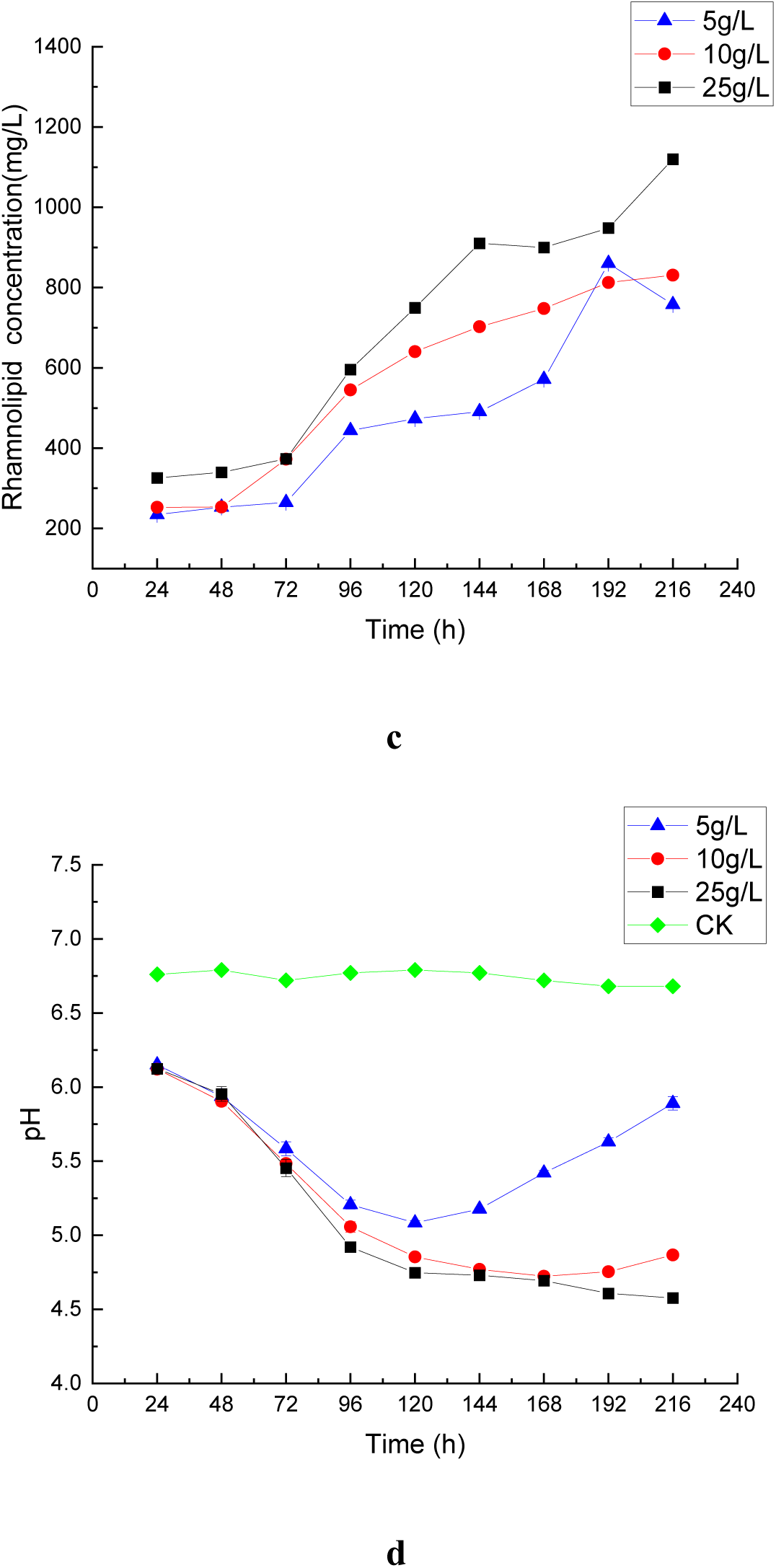
Time course profiles of **a** cell density, **b** lipase activity, **c** rhamnolipid concentration, and **d** pH by *P. aeruginosa* M4 degraded different concentrations WCO.

When fermentation was carried out using an initial WCO concentration of 5 g/L, it was found that the pH decreased with the accumulation of medium-long-chain fatty acids before 120 h. After 120 h, the pH increased gradually. However, during this period, cell concentration continued to decrease and rhamnolipid concentration continued to increase. Thus, it could be concluded that rhamnolipid was being produced as a secondary metabolite.

### Salt tolerance of *P. aeruginosa* M4

Due to the high salt content in WCO, it is necessary to assess the salt tolerance of the isolated strain. The growth of *P. aeruginosa* M4 in media with varying salt concentration is shown in Table 4. Here it could grow well up to 70 g/L NaCl, while the *Pseudomonas* selected by Li et al. (2001) could grow at a salt concentration of between 10 g/L and 50 g/L. When NaCl concentration reached between 100 g/L and 150 g/L, the growth of *P. aeruginosa* M4 was significantly inhibited, and the optimal cell density was observed at a salt concentration of 10 g/L. The ability of *P. aeruginosa* M4 to grow at higher salt concentration up to 70g/L suggests that it may be an ideal candidate for applications using high salt of WCO as the primary carbon source.

**Table 4.** Salt tolerance of *P. aeruginosa* M4.

| Salt concentration(g/L) | 0 | 2 | 5 | 10 | 15 | 20 |
| --- | --- | --- | --- | --- | --- | --- |
| Cell density | 1.33±0.05 | 1.37±0.05 | 1.47±0.02 | 1.49±0.01 | 1.44±0.01 | 1.36±0.13 |
| Salt concentration(g/L) | 25 | 30 | 50 | 70 | 100 | 150 |
| Cell density | 1.32±0.06 | 1.24±0.04 | 1.13±0.02 | 0.78±0.02 | 0.04±0.01 | 0.03±0.01 |

## Discussion

Most households in China like to use a large amount of edible oil for cooking dishes, so the output of WCO is huge (Demichelis et al. 2017). At present, there is no effective way for WCO treatment and utilization, which results in various environmental and social problems (Zhang et al. 2010). Funston et al. (2016) and Pacheco et al. (2012) studied the rhamnolipid production using glycerol as a substrate, and Müller et al. (2010) reported the rhamnolipid production with sunflower oil. However, there were few reports on the rhamnolipid production from WCO. The rhamnolipid production from WCO is expected to be a win-win method for both effectively treating WCO and economically producing biosurfactants. The *P. aeruginosa* M4 isolated in this study had a rhamnolipid production capacity up to 1119.87 mg/L, which was much higher than the reported rhamnolipid concentrations of 430 and 299 mg/L produced by *P. aeruginosa* (Radzuan et al. 2017; Ramírez et al. 2016). Moreover, it could tolerate a high concentration of salt, which usually was contained in WCO. This suggested that *P. aeruginosa* M4 may be a good candidate for both WCO treatment and biosurfactants production.

Various types of carbon sources including glucose, fructose, malt extract powder, and soluble starch were usually used as the substrate for lipase production (Zamroni 2009). However, the carbon catabolic regulation in microbial systems preferentially catabolizes the best carbon source, the one which facilitates cell growth being used before any of the others (Saxena et al. 2003). It is well known that organic nitrogen sources can affect enzyme synthesis because they provide the amino acids and other growth factors required for cell metabolism and protein synthesis (Iftikhar et al. 2011). This study showed that oil and tryptone was the best carbon and nitrogen source to maximize the bacteria growth and lipase activity. Low inoculation ratios could prolong the lag phase of cell growth, while high inoculation ratios might lead to restrictions in nutrient and oxygen availability. The optimum inoculum ratio of 5% (v/v) was obtained in this study. Temperature affects the secretion of extracellular enzymes. The temperature of 35°C achieved the maximum lipase activity (19.87 U/mL), which was consistent with the result reported by Prasad (2014). The pH of the environment greatly affects cell growth and lipase activity. Usually, the acidic pH is most suitable for enzyme production (Hiol et al. 2000). Diaz et al. (2006) reported that the optimal pH was 6.5 for lipase production by *Rhizopus homothallicus*, while it was 8.0 in this study. This may be because that the higher initial pH could enable the fermentative system to tolerate higher concentrations of fatty acids.

Combined with previous study, the formation pathway of rhamnolipid was illustrated in Fig. 6. Abdel-Mawgoud et al. (2011) and Gutiérrez-Gómez et al. (2019) described the metabolic pathway of rhamnolipid biosynthesis from glucose, where both R-3-hydroxyalkanoate and the rhamnose precursor were synthesized from glucose. In this study, the only carbon source for rhamnolipid production was WCO. In case of WCO, the rhamnose precursor could be synthesized from glycerol, which was a hydrolysis product of WCO. Theoretically, the C10 medium-long chain fatty acids of oil degradation products might be directly used as R-3-hydroxyalkanoate precursor. However, Abdel-Mawgoud et al. (2014) pointed out that, R-3-hydroxyalkanoate precursor was synthesized de novo from acetyl-CoA, even with fatty acids as a carbon source. The C10 medium-long chain fatty acids could not be directly transformed to the R-3-hydroxyalkanoate precursor.

**Fig. 6.**
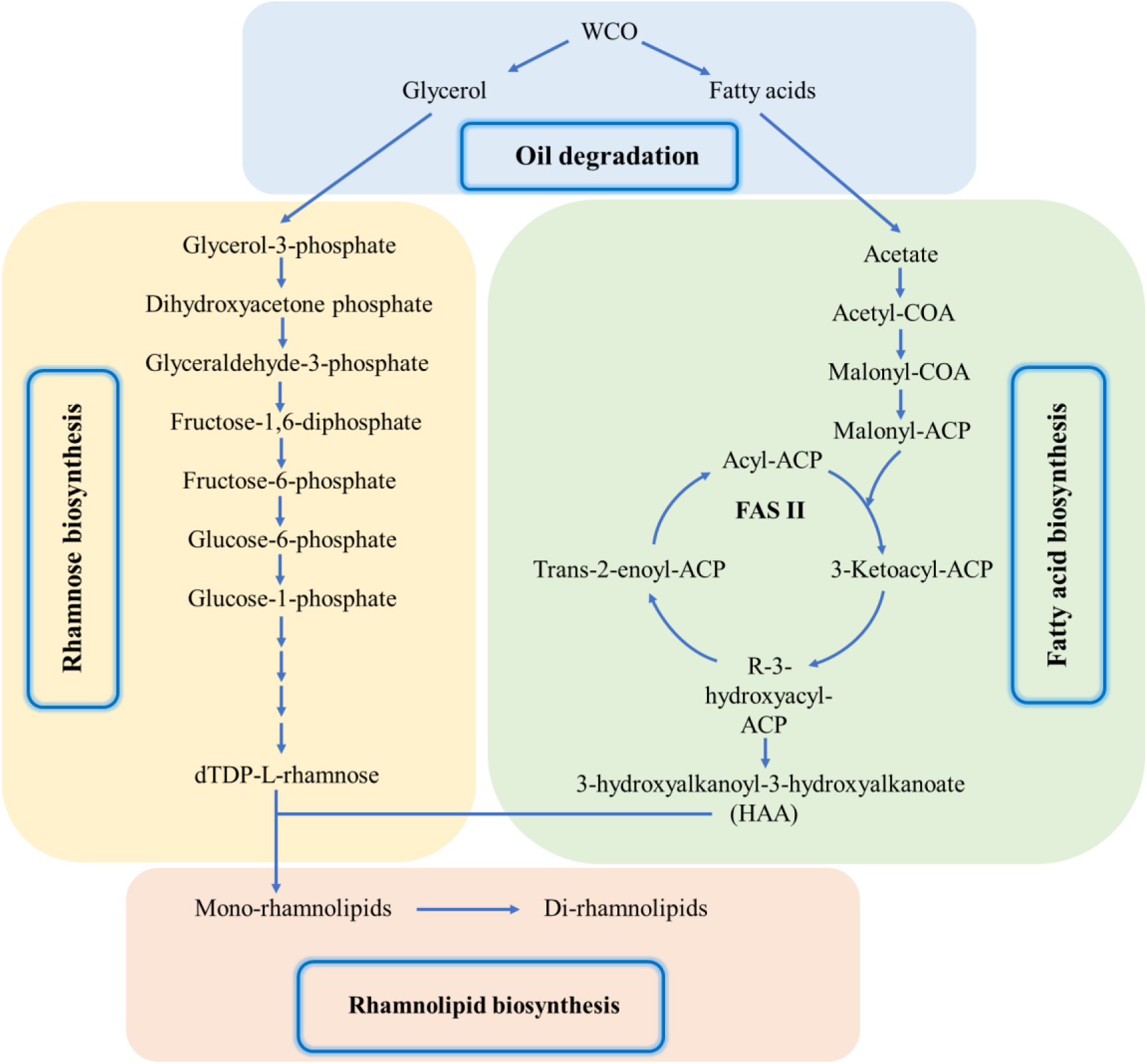
Proposed synthetic pathway of rhamnolipid using WCO as sole carbon source.

## Author contributions

JS, XFL and DL conceived and designed the research. JS and YCC performed the experiments. JS, YR and DL analyzed the date. JS and DL wrote the manuscript. All authors read and approved the manuscript.

## Compliance with ethical standards

### Ethical approval

This paper does not contain any studies with human participants or animals.

